# Targeted induction of gut-microbial metabolism acutely affects feeding patterns and clock gene expression in the host

**DOI:** 10.1101/2023.06.20.545777

**Authors:** Giorgia Greter, Claudia Moresi, Stefanie Oswald, Alice de Wouters d’Oplinter, Daria Künzli, Elisa Cappio Barazzone, Jiayi Lan, Emma Slack, Markus Arnoldini

**Affiliations:** Department of Health Sciences and Technology, ETH Zurich, 8093, Zurich, Switzerland; Department of Chemistry and Applied Biosciences, ETH Zurich, 8093, Zurich, Switzerland

**Keywords:** Gut microbiota, diurnal rhythm, microbial metabolism, fermentation products

## Abstract

The gut microbiota and host diurnal rhythm mutually influence each other, and microbiota metabolism has been shown to play a role in regulating host circadian function via secretion of fermentation products. Microbial metabolism is dependent on the availability of nutrients for the microbiota, typically through the host’s food intake, making it challenging to disentangle the effect of host and microbiota metabolism. In this study, we acutely induced gut microbial metabolic activity without inducing host metabolism in mice. We found that increasing microbial metabolism in the gut altered clock gene expression locally. Actuating microbiota metabolism also reduced host food intake beyond the calories provided by the microbiota, suggesting a systemic signaling effect of microbial metabolism on the host.

## Introduction

From regulating our daily sleep-wake cycles to influencing metabolic processes, the shift between day and night dominates the rhythm of most life on Earth (*1*, *2*). The mammalian circadian clock is a fundamental biological process which functions through a tightly regulated network of transcriptional feedback loops (*3–7*) that trigger periodic fluctuations in physiology and behavior (*8*, *9*). One of the key behavioral rhythms that is regulated by the circadian clock is food intake (*10*). In anticipation of the changing food availability with a change in light, the master pacemaker activates orexigenic pathways and hormone production to increase feeding and activity, which in turn triggers anorexigenic pathways through satiety signals to prepare for fasting and rest (*11*). Synchronization of these behaviors with the natural light-dark cycle increases fitness, while being asynchronous with the external environment is detrimental to health (*12–14*).

Recent research has revealed crucial links between the biological clock, feeding behavior, and the intricate ecosystem of the gut microbiota (*15*, *16*). The circadian oscillations in food intake in the host result in oscillations in gut microbial metabolism and composition in a circadian pattern (*17*, *18*). Oscillating immune function in the host also plays a role in shaping the gut microbial ecosystem. For example, immunoglobulin A, a key antibody that regulates commensal bacteria community composition, is secreted with a circadian rhythm, and host responses to the invasion of pathogenic bacteria are dependent on the time of day (*19–21*).

In addition, there is evidence that this regulation goes both ways: not only does the host clock regulate the microbiota, but also vice versa: the microbiota plays a significant role in the amplitude of host clock gene expression oscillations, and is associated with weight gain and reduced longevity in a dysregulated circadian rhythm (*22–25*). In line with this, germ-free (GF) mice, which do not have a microbiota, are less sensitive to the dysregulation of the circadian clock induced by high fat diets (*26*).

A major way of signaling between the gut microbiota and the host are microbial fermentation products (*27*). Fermentation products are metabolic byproducts of the microbiota which are used by the host for many metabolic and endocrine functions in the gut-brain axis, as well as for food intake regulation (*28–30*). Fermentation product excretion by the microbiota can trigger satiety signaling in the host and may play a role in regulating the host circadian rhythm due to the circadian nature of food intake (*17*, *31*). However, metabolic oscillations in the gut microbiota arise from host food intake, making it difficult to disentangle host effects from microbial effects, and thus to evaluate mechanistic interactions between the microbiota and the host circadian clock (*32*).

In this study, we use an treatment with lactulose, a disaccharide that is not taken up in the small intestine of mice but can be metabolized by the gut microbiota, to increase microbial metabolism in the gut without directly providing calories to the host. This changes the normal microbial rhythm that depends on food intake patterns of the host The effect of this intervention is transient, and the system recovers rapidly from this disruption without lasting changes in the diurnal rhythm after an acute disruption. This experimental setup allowed us to evaluate how microbiota activity regulates host circadian clock gene expression and subsequent eating patterns. We show that microbial metabolism influences host clock gene expression and that this systemically regulates host feeding behavior.

## Results

### Microbial metabolism fluctuates with the host’s diurnal feeding pattern

We first asked whether gut microbes exhibit diurnal variation in population size in a defined setting. To address this, we quantified bacterial load in fecal samples collected every 6 hours over two consecutive days. For these experiments, we used gnotobiotic mouse model colonized with a minimal microbiota of three species (3MM), consisting of *Bacteroides thetaiotaomicron* (*B. theta*)*, Eubacterium rectale* (*E. rectale*), and *Escherichia coli* (*E. coli*), maintained across multiple generations. These species represent the phyla Bacteroidetes, Firmicutes, and Proteobacteria (recently renamed to Bacteroidota, Bacillota, and Pseudomonadota), respectively, which together dominate the gut microbiota (*15*, *33*), and all three species reach high numbers in the gut of colonized mice (Figure 1A). Despite prior observations of phylum-level oscillations associated with feeding cycles (*34*), we detected no significant rhythmic changes in the population sizes of any of the three species over the course of two 24 hour cycles (Figure S1A).

**Figure 1.**
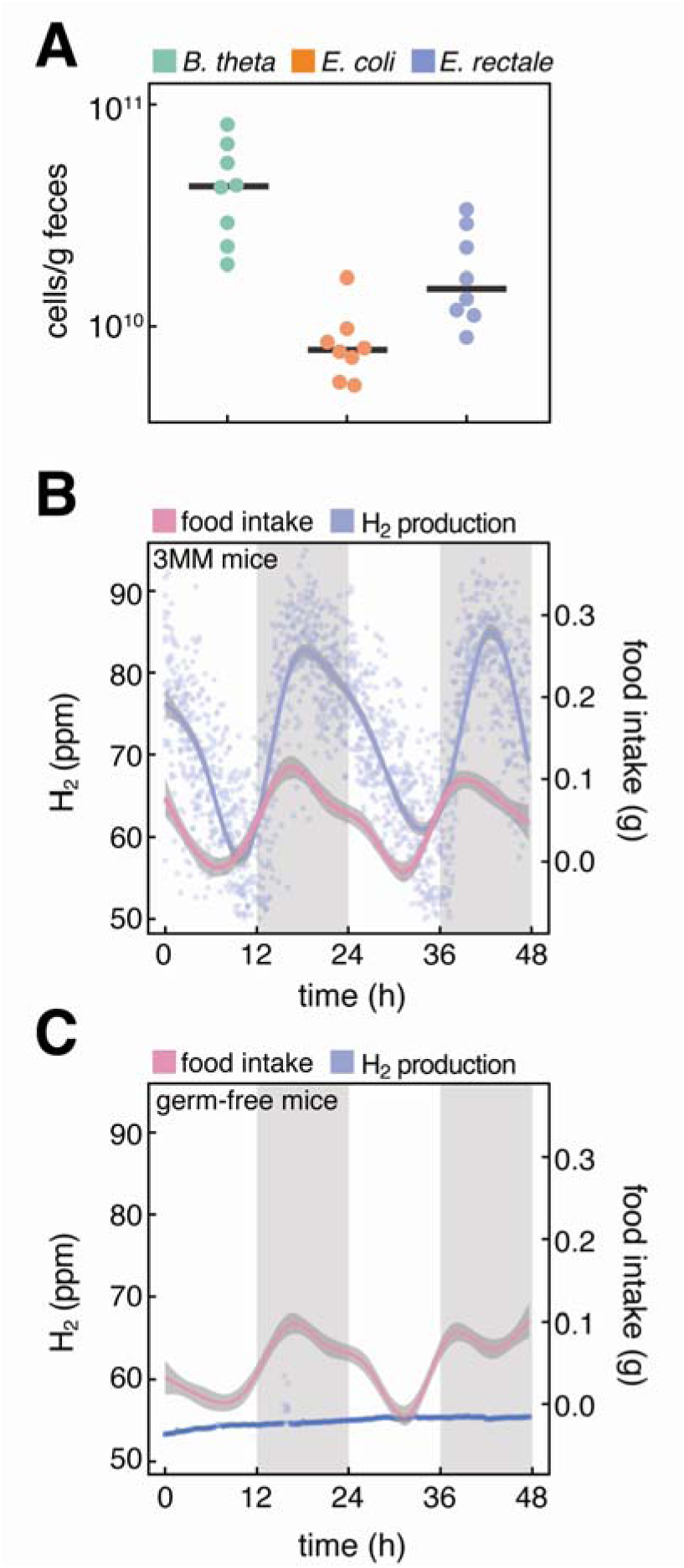
Timing of host food intake and microbial activity are linked. **(A)** Cell counts per gram feces in the EAM mouse (n=8) at the end of the light cycle (zt=12h), measured by qPCR. **(B)** Food intake and hydrogen production of the EAM mouse over time. Two days of measurement are shown, measurements are taken every 24min (n=16). Curves were calculated using the geom_smooth function of the package ggplot2 in R with standard settings. SE is illustrated with grey shading surrounding the trend line. **(C)** Food intake and hydrogen production of the germ-free mouse over time (n=8).

Food intake was measured for each mouse using a TSE Phenomaster system that is fully enclosed in an experimental isolator. We have previously developed this system, which included extensive testing for sterility or gnotobiosis (*35*). As a readout for microbial metabolism, hydrogen gas excretion was also measured over time (Figure 1B and Figure S1B). Hydrogen is an abundant byproduct of butyrate production, and *E. rectale* is the only butyrate producer in the 3MM, and therefore the dominant hydrogen producer (Figure S1C) (*36*, *37*). Hydrogen levels fluctuated in a 24-hour period in the 3MM mouse, closely following mouse food intake, which provides the non-host-available fibers that the microbiota can metabolize in the gut (Figure 1B, Figure S1B). Germ-free mice did not produce measurable levels of hydrogen despite having the same food intake rhythm (Figure 1C).

### Lactulose treatment acutely disrupts the diurnal pattern of microbial metabolism

To acutely disrupt the normal diurnal rhythm of microbial metabolism in the gut without directly providing nutrients to the host, we administered lactulose to 3MM mice. Lactulose is a disaccharide that cannot be metabolized by mammals, is known to be metabolized by the gut microbiota, and is associated with an increase in hydrogen production and reduced colon pH due to microbial metabolic activity (*38–40*). We administered the lactulose between Zeitgeber times 3 and 4, several hours after the mice had stopped eating and when hydrogen production was decreasing. To control for a potential effect of the oral gavage itself on the microbiota, PBS was administered as a negative control, and all experiments were replicated in germ-free mice to control for host effects independently of the microbiota.

Mice that were treated with lactulose showed increased hydrogen production two hours after treatment compared to PBS-treated mice, which continued in the normal rhythm of hydrogen production (Figure 2A). There was a significant difference in measured hydrogen levels between the mice in the two groups at the endpoint of the experiment at zt=8, 5h after lactulose treatment, indicating that the microbiota had metabolized the administered lactulose (Figure 2A, right). To evaluate the duration of the lactulose-dependent change in the diurnal pattern of microbiota metabolism, we repeated the experiment and monitored hydrogen production over the following dark cycle. Hydrogen levels in lactulose treated mice were equal to PBS treated mice by Zeitgeber 15, indicating that the microbiota had metabolized the lactulose and resumed its normal metabolic rhythm, and that our treatment therefore constitutes an acute and transient intervention (Figure 2B). To test whether these results were translatable to a complex microbial community, we performed this experiment in Specific Pathogen Free (SPF) mice. This effect was equally observed in SPF mice, indicating that a complex microbial community also metabolizes lactulose in an acute manner (Figure S2A).

**Figure 2.**
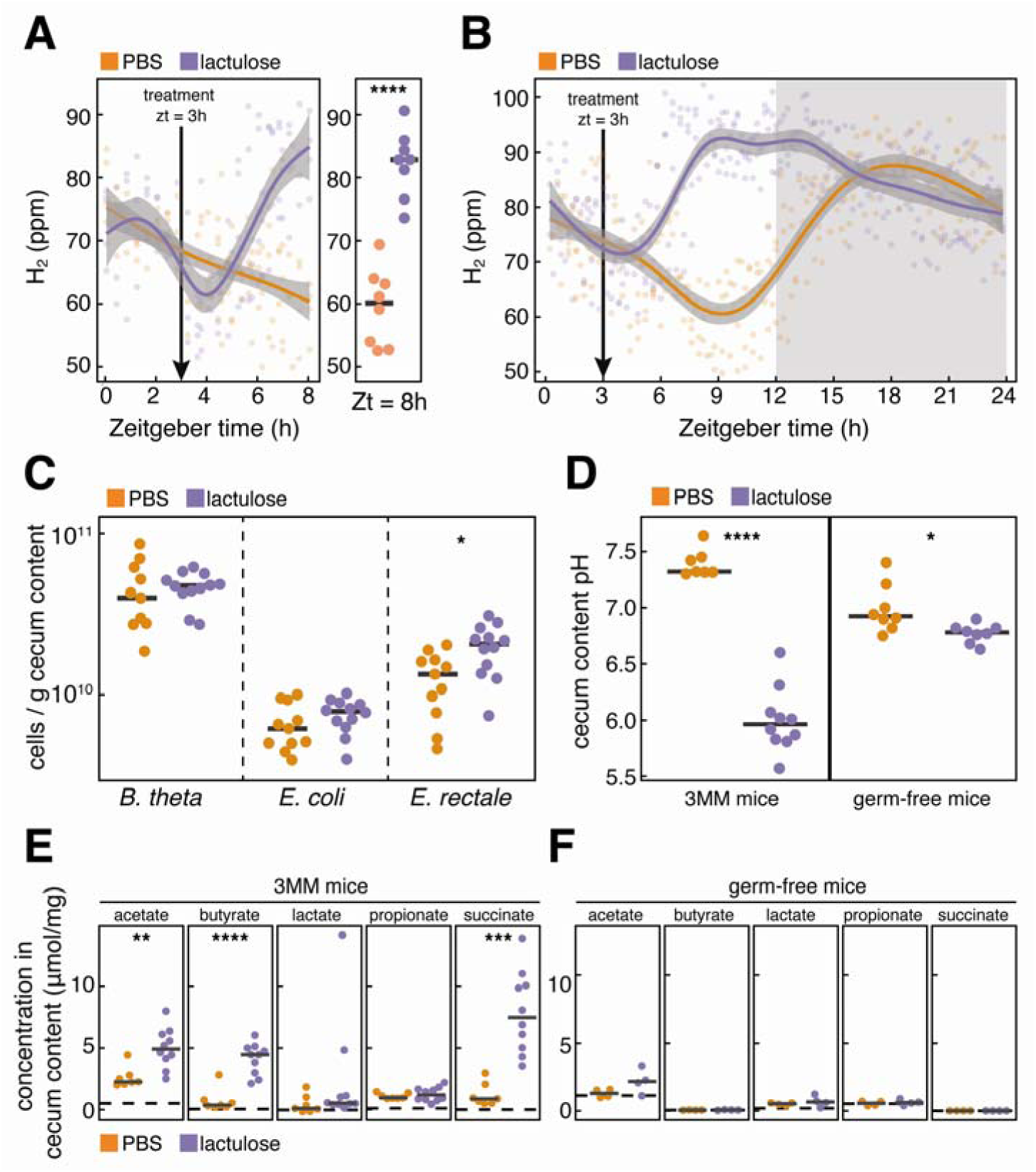
Lactulose treatment disturbs the diurnal rhythm of microbial metabolism. **(A)** Hydrogen production increases after lactulose treatment (purple, n=8), but not after treatment with PBS (orange, n=8). Arrows at zt=3h indicate treatment. An unpaired t-test was performed on final hydrogen measurements (p>0.0001). **(B)** Hydrogen production after lactulose treatment (purple, n=6) is temporary and reverses back to normal levels after treatment, and is not observed in PBS treatment (orange, n=5). **(C)** Bacterial counts in cecum 5h after lactulose (purple, n=12) and PBS treatment (orange, n=10). *E. rectale* population increases in lactulose treated mice compared to PBS treated mice (p<0.05), *B. theta* and *E. coli* counts do not change significantly. **(D)** Cecum pH of 3MM (n=10 for lactulose, n=7 for PBS treatment) and germ-free (n=8 in each group) mice 5h after treatment. The pH decreases after lactulose treatment in 3MM (p<0.0001, unpaired t-test) and germ-free (p<0.05) mice. **(E)** Concentrations of microbial metabolites in cecum content of 3MM mice 5h after treatment with lactulose (purple, n=10) or PBS (orange, n=7). Acetate, butyrate, and succinate concentrations increase in lactulose treated as compared to PBS treated mice (**, ***, **** indicate significance levels smaller than 0.01, 0.001, and 0.0001, respectively); unpaired t-test, Benjamini-Hochberg corrected for multiple testing. **(F)** Concentrations of microbial metabolites in cecum content of germ-free mice 5h after treatment with lactulose (purple, n=8) or PBS (orange, n=8). There is no significant difference between groups (unpaired t-test, Benjamini-Hochberg corrected for multiple testing).

Five hours after the initial lactulose treatment, after the microbiota had metabolized the lactulose, the mice were euthanized, and bacterial population sizes in cecum content were measured to evaluate the effect of lactulose on the 3MM microbiota. There was a significant increase in the cecal *E. rectale* population, but no significant change in the numbers of *B. theta* or *E. coli* (Figure 2C). Having observed increased hydrogen production in lactulose-treated mice, we further evaluated the effect of lactulose on microbial metabolism. Microbial metabolism results in an increase in excretion of acidic fermentation products, which should lead to a decrease in pH in the cecum. There was a significant drop in cecum content pH when 3MM mice were treated with lactulose, which was associated with an increase in fermentation product concentration in the cecum (Figure 2D and 2E). Lactulose-treated mice had significantly higher concentrations of acetate, butyrate, and succinate, indicating increased metabolic activity of all 3MM species, despite the lack of significant increase in *B. theta* and *E. coli* population densities (*41*, *42*). *E. rectale* is the principal butyrate producer in the 3MM, while *B. theta* and *E. coli* both produce acetate, and *E. coli* produces succinate when grown on lactulose (Figure S2B). Germ-free mice that were treated with lactulose showed a small but significant drop in pH in cecum content, likely due to osmotic effects (Figure 2D). In GF mice, fermentation product concentrations did not increase after lactulose treatment (Figure 2F).

To ensure that the observed effects of lactulose treatment are due to the induction of microbial metabolism and not osmotic effects of lactulose treatment in our mouse models, we measured the water content of feces and cecum content after treatment in 3MM mice and germ-free mice in all experiments. In contrast to what would be expected for an osmotic laxative, water content in fresh feces of 3MM mice and the total fecal output in the 5h following treatment decreased with lactulose treatment, possibly due to the increased microbial biomass from lactulose metabolism, and indicating that the osmotic effect of lactulose treatment was small in these mice (Figure S2C,D). The water content of cecum content in 3MM and germ-free mice (Figure S2E) did not differ significantly between the different treatments. A factor that could potentially influence these parameters is the difference in cecum sizes between germ-free and colonized mice. While the cecum of 3MM mice is significantly smaller than that of germ-free mice (Figure S2F), both mouse models have large ceca when compared to SPF mice(*35*). Following up the surprising lack of an osmotic effect of lactulose treatment in 3MM mice, we treated 3MM and SPF mice with twice the amount of lactulose and measured fecal water content over time (Figure S3A), water content of cecum content (Figure S3B) and total fecal output during 5h after treatment (Figure S3C). Interestingly, lactulose only had a significant effect on the water content of fresh feces and cecum content in SPF mice, but not in 3MM mice. We concluded that the osmotic effect of lactulose is dependent on the colonization state of the animals, and not a major contributor to the observed effects in 3MM mice.

### Inducing microbial metabolism acutely increases clock gene expression in small intestinal tissue

To test whether the induction of microbial metabolism affects gene expression driving the host clock, we studied the transcription of several core clock genes in the mouse gastrointestinal tract (Figure 3A). Clock regulation begins with the core clock genes (*clock* and *arntl),* which activate transcription of the *per* and *cry* repressor genes (*43*, *44*). *Clock* and *arntl* also activate transcription of the *rev-erb* and *ror* genes in a second regulatory loop. Finally, *clock* and *arntl* regulate the transcription of *nfil3* and *dbp* in a third regulatory loop. These genes all function in a transcriptional autoregulatory feedback loop throughout the body.

**Figure 3.**
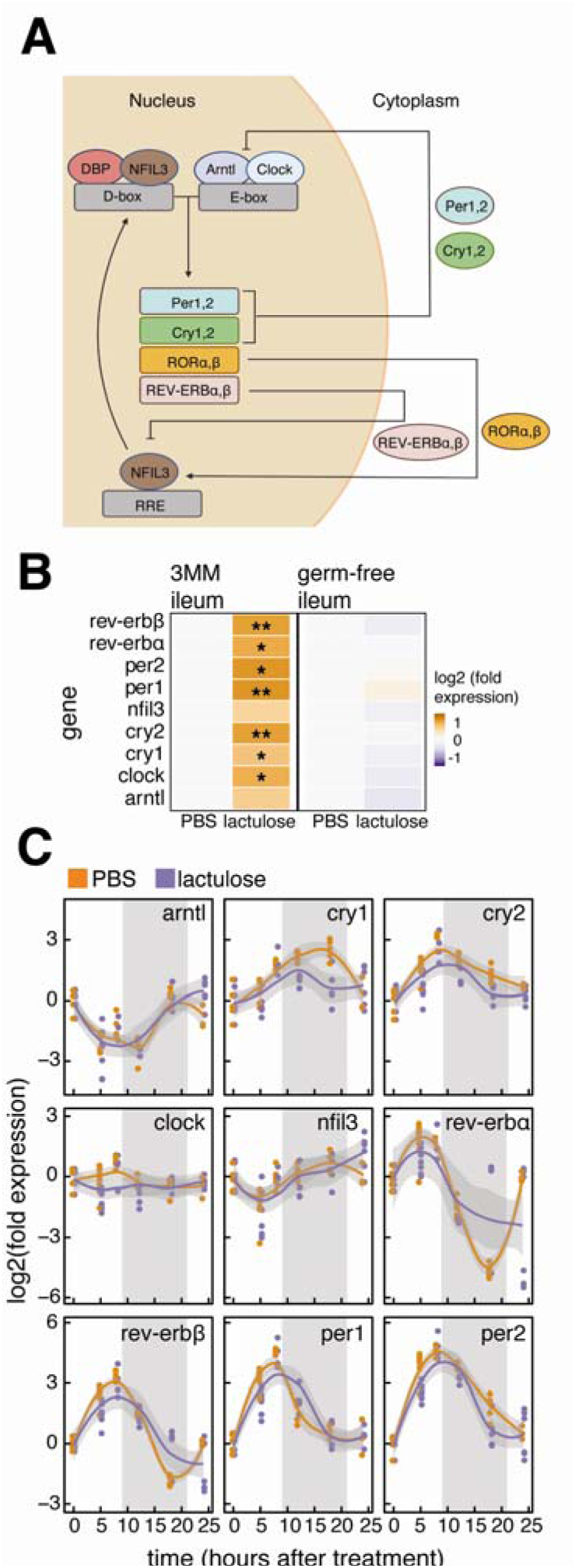
Lactulose feeding acutely alters clock gene expression in small intestinal tissue. **(A)** Schematic of the clock gene network in mice. **(B)** Clock gene expression in ileum tissue in 3MM (N=10 for lactulose treatment, n=7 for PBS treatment) and germ-free (n=8 in each group) mice 5h after treatment, measured via qPCR. Fold change values are relative to the average of the respective PBS control groups. Statistical significance was tested using upaired t-tests and Benjamini-Hochberg corrected for multiple testing (* and ** indicate p<0.05 and p<0.01, respectively). **(C)** Clock gene expression in ileum tissue of 3MM mice during 24h after treatment (treatment starts at time 0h). All gene expression data was normalized to samples collected from untreated mice at zt3 (control). Expression differed significantly between treatment and control groups for cry1, cry2, and per2 (two-way ANOVA (time, treatment), p<0.05).

As described above, 3MM mice were treated with lactulose or PBS between zeitgeber time 3-4 and euthanized five hours later. Tissue was harvested from the ileum, cecum, and colon for RNA extraction and RT-PCR. The gene expression of core clock genes was compared between the two treatment groups. When 3MM mice were treated with lactulose, there was a significant increase in several core clock genes in the ileum tissue, but no significant difference in gene expression in cecum and colon tissues, where most microbial metabolism occurs (Figure 3B and S3A). These differences in expression reflected the fluctuations seen in peripheral tissues in the normal mouse circadian rhythms (*45*). To control for the direct effect that lactulose may have on the host, this experiment was also conducted in germ-free mice. When germ-free mice were treated with lactulose, there was no significant increase in core clock genes in any of the studied tissues, indicating that microbial metabolism induced changes in the peripheral circadian clock (Figure 3B and S3B). Furthermore, to test whether this effect was translatable to a complex microbiota, this experiment was conducted in SPF mice. When SPF mice were treated with lactulose, there was a significant decrease in *cry1* expression in the ileum, indicating that the effect of microbial metabolism on SPF clock gene expression was diminished (Figure S3C).

To evaluate the extent of the effect of acute microbial metabolism on clock gene expression, we measured expression in ileum tissue at several time points over 24 hours following lactulose or PBS treatment in 3MM mice (Figure 3C). We found that gene expression of *cry1, cry2,* and *per2* was significantly different after treatment. This finding indicates that the effect of lactulose on microbial metabolism also resulted in an effect on host clock gene expression.

### Inducing microbial metabolism decreases host food intake in following active cycle

We then tested whether increasing host clock gene expression through specific alteration of gut microbial metabolism has further effects on the behavioral rhythm of the host. We measured the rate and total amount of 3MM mouse food intake in the dark cycles before and after lactulose treatment. The dark cycle preceding treatment was used to ensure normal feeding behavior for each mouse and provided an undisturbed baseline measurement. Lactulose or PBS treatments were then administered as before at zeitgeber 3-4, and food intake was measured again in the following dark cycle and compared to the baseline food intake in the previous dark cycle. 3MM mice treated with lactulose reduced both their food intake rate and total food intake in the following active cycle, while there was no significant reduction in food intake in 3MM mice that were treated with PBS (Figure 4A and S5A). We subsequently continued to measure food intake in 5 lactulose-treated mice and 5 PBS-treated mice for a second active cycle, and found that the decrease in food intake did not persist, indicating an acute effect of microbial metabolism on food intake (Figure S5B). When repeating this experiment with treatment during the active phase at zt=14-15, we did not see a significant change in food intake in the dark cycle after treatment (Figure S5C, D).

**Figure 4.**
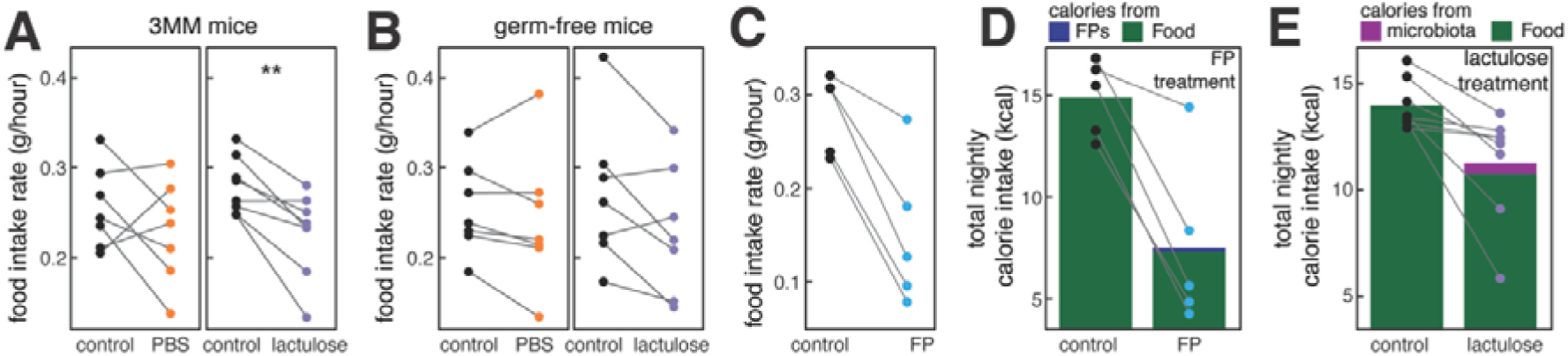
Lactulose treatment decreases host food intake in the following active cycle. **(A)** Food intake rate in 3MM mice in the dark phase prior to treatment (control) and following treatment (treatment), after lactulose (n=8) and PBS (n=7) treatment. Food intake is significantly reduced after lactulose treatment (paired t-test, p<0.01). **(B)** Food intake rate in germ-free mice after lactulose (n=5) and PBS (n=4) treatment. Food intake is not significantly different between groups (paired t-test). **(C)** Food intake rate of 3MM mice during the dark phase prior to treatment with fermentation products (control) and the dark phase after treatment (treatment). Food intake is significantly reduced after treatment (paired t-test, p<0.01). Treatment was done by oral gavage of 66mg of fermentation product in 100µL of PBS, with a 4:3:3 ratio of sodium succinate, sodium acetate, and sodium butyrate, respectively. **(D)** 3MM mouse total calorie intake after accounting for the calories directly provided by fermentation products in treatment. Calory intake remains significantly reduced after treatment when calories provided by the treatment are considered (paired t-test, p<0.05). **(E)** 3MM mouse total calorie intake after accounting for calories provided by the microbiota after lactulose metabolism. Calory intake remains significantly reduced after lactulose treatment when calories provided by microbial metabolism are taken into account (paired t-test, p<0.05).

We repeated this experiment in GF mice and SPF mice and did not find any significant reduction in mice treated with lactulose (Figure 4B and S5E, F). The non-significant trend towards lower food intake in lactulose-treated GF mice can potentially be explained by the viscosity of the lactulose solution; viscous solutions given to mice via oral gavage have been shown to decrease food intake (*46*). However, the decrease in food intake is small in GF mice compared to 3MM mice, indicating that actuation of microbiota metabolism is the dominant cause for this of this effect.

To evaluate the mechanism by which microbial metabolism decreases food intake in 3MM mice, we allowed 3h for gene expression changes to take place in peripheral tissues after the peak of bacterial activity at 5h after lactulose administration. We thus measured circadian clock gene expression in the 3MM hypothalamus and liver 8h after lactulose or PBS treatment (Figure S6A). We did not find any significant change in clock gene expression in either of these organs, indicating that the mechanism does not lie in a systemic shift in circadian clock gene expression in hypothalamus or liver, at least at the timepoint we investigated.

### Disruption of food intake patterns is due to systemic effects of microbial metabolic products

Microbial metabolism of lactulose in the mouse takes place largely in the cecum and colon (*47*). However, core clock gene expression was most significantly altered in the ileum when mice were administered lactulose. These mice then consumed less food following treatment, implying that the effect of microbial metabolism on the host is systemic and not localized to the point of metabolic processes (Figure 3B and 4A). To test the systemic effect of microbial metabolism on host food intake, 3MM mice were administered a fermentation product mix of succinate, acetate, and butyrate, as these were the fermentation products that were significantly produced by the 3MM in the cecum after lactulose treatment (Figure S2B). Orally administered fermentation products are taken up by the host in the small intestine, before reaching the cecum (*48*). 3MM mice reduced their rate of food intake and their total food intake following the treatment of fermentation products, indicating that the effect of the microbiota on host circadian behavior is systemic (Figure 4C). When we repeated this experiment in SPF mice, we did not find a significant decrease in food intake following fermentation product treatment (Figure S6B). This is possibly due to uptake efficiency differences in the SPF small intestine compared to the 3MM mouse, or due to differences in metabolic signaling processes between these mice.

Providing calories in the form of fermentation products is one potential mechanism by which the microbiota regulates host food intake systemically (*49*). To evaluate whether the reduction in host food intake was solely due to the supplemental energy provided by the fermentation products, we estimated the total number of kilocalories that were taken up by the host before and after the treatment with fermentation products, assuming that all orally administered fermentation products serve as an energy source (0.19 kcal). We found that mice reduced their food intake by more than these 0.19 kcal, going beyond compensation for the additional energy provided by the microbiota (Figure 4D).

To test if this effect of the fermentation products could be observed when they were not administered orally but produced by microbial metabolism, we estimated the calories in fermentation products produced by the microbiota after lactulose treatment (Figure 4B). To get an upper bound for this number, we assumed that the energy in fermentation products has to be lower than the combustion enthalpy of the fed lactulose (0.52 kcal). We found that 3MM mice reduced their food intake following lactulose treatment by even more than these 0.52 kcal, showing that microbiota metabolism affects host behavior through a mechanism other than energy homeostasis of the host (Figure 4E).

To understand the mechanism by which microbial metabolism regulates host food intake, we measured various satiety-associated hormones in 3MM, SPF, and GF mice. We measured concentrations of the hormones PYY and GLP-1, known to be associated with gut microbiota changes (*50*, *51*), in portal vein plasma 5h after treatment with PBS or lactulose. We found no significant difference in PYY or GLP-1 concentrations between control and treatment groups in 3MM, GF, or SPF mice 5h after treatment (Figure S7A, B). We also measured ghrelin and leptin concentration in cardiac serum of 3MM and GF mice 5h after PBS or lactulose treatment and found no significant effect of lactulose on their concentrations in 3MM mice 5h after treatment (Figure S7C, D). There was a small but significant increase of ghrelin concentration in lactulose treated GF mice, potentially due to the osmotic effect of lactulose increasing hunger signaling (Figure S7C).

## Discussion

The gut microbiome and host circadian clock are intricately linked, with evidence suggesting that the host circadian clock can influence the gut microbiota and vice versa (*34*, *52*). In this study, we show that the microbiota regulates host circadian clock gene expression and feeding behavior through metabolic rhythmicity. We first demonstrated that the EAM gut microbiota oscillates metabolically in a periodic manner which is closely associated with the periodic food intake of the host. Similar rhythmic oscillations in bacterial metabolism have been found independently of microbiota complexities (*18*, *35*, *53*). We disrupted the metabolic rhythmicity of the 3MM by providing a nutrient that is available only to the microbiota during the host’s inactive period, during which food intake is typically low (Figure 1C). We observed an acute increase of microbial metabolism following one treatment of lactulose that was comparable to normal production during the host’s active state, simulating host food intake for the microbiota (*17*). Consequently, we observed an increase of expression of several clock genes in the ileum tissue, some of which persisted for several hours (*45*).

Most microbial metabolism takes place in the cecum, and interestingly, we found no change in cecum tissue gene expression after treating 3MM mice with lactulose. This may indicate a systemic effect of microbial metabolism on the host, rather than a localized one. In fact, a systemic relationship between the gut microbiota and small intestinal epithelial rhythmicity has previously been described as it was found that altering the microbiota through diet causes changes in small intestinal immune function (*54*, *55*). The systemic effect of the microbiota on the host clock gene expression likely influenced host food intake following the lactulose treatment. We observed an increase in fermentation product concentration in the mouse cecum after administering lactulose to 3MM mice, and a subsequent decrease in food intake in the following active cycle. We observed a similar decrease in food intake when we administered a mix of fermentation product directly to the host, indicating that microbial metabolism is in part regulating host feeding behavior. One possible mechanism by which this may occur is through microbial feeding of the host (*49*). The microbiota regulates host appetite through energy harvesting of inaccessible fibers: GF mice eat more food in a day than colonized mice to compensate for the reduced energy extraction in the large intestine that is otherwise provided by the microbiota (*35*). By treating 3MM mice with lactulose, we assumed that the host would receive energy indirectly from microbial fermentation, and that the host would reduce its food intake in the following dark cycle by the same number of kilocalories. However, after accounting for the energy that the microbiota was providing the host through lactulose metabolism, we still observed a reduction in total energy intake in mice. We observed similar results when we provided the host directly with fermentation products. This finding indicates that providing energy is not the only mechanism by which microbial metabolism reduces host feeding, and that there is a further mechanism that regulates the host’s decrease in total energy harvest.

Surprisingly, when we repeated our experiments in SPF mice we did not observe the same results as in 3MM mice. First, we did not measure the same changes in clock gene expression in mice that were treated with lactulose. Furthermore, mice treated with lactulose did not decrease their food intake in the following active cycle. We hypothesized that the microbial composition of the SPF mice resulted in the utilization of fermentation product that were generated after lactulose treatment. However, direct treatment of SPF mice with fermentation product also did not decrease their food intake, indicating that this was not the case. One possible explanation for this difference might be that SPF mice have decreased sensitivity to changes in fermentation products, and that further lactulose doses would induce similar changes in clock gene expression and food intake in SPF mice. Alternatively, there might be a signaling factor in the SPF microbiota that is absent in the 3MM, which can block the systemic effect of fermentation products downstream of uptake into the systemic circulation.

The experimental approach of using lactulose feeding to acutely induce microbial metabolism also has its limitations. Lactulose can have an osmotic effect, which we found to be dependent on microbiota complexity (Figure S3), and could possibly also have microbiota independent effects on the host. While we tried to control for these effects by ensuring that the osmotic effect in our sytem was minimal (Figure S2C-E), it would be desirable to use different methods of specifically actuating microbiota metabolism in future studies. One possibility could be the use of genetic systems that can be activated by experimentally tunable chemical inducers (*56*), which may avoid the more complex physiology of lactulose metabolism.

Administration of fermentation products may have a systemic effect on the host through various mechanisms, and there is an association between fermentation product intake and a reduction in food intake (*51*, *57–59*). Butyrate dietary supplementation in mice decreases food intake and promotes fat oxidation, and acetate supplementation can decrease appetite directly (*60*, *61*). It has also been found that dietary supplementation of fermentable carbohydrates which can be broken down by the microbiota result in changes in regulation of satiety hormones which are produced in a circadian manner (*50*, *62–64*). However, we did not find any of these hormones to be acutely altered 5h after treatment of mice with lactulose, indicating that the systemic effect of microbial metabolism on food intake might involve a different mechanism. However, it remains possible that changes in hormone levels happen at a different time point, and a systematic analysis of microbial effects on the temporal development of hormone levels would be of intertest. It has previously been found that mice with gene deletions in the circadian clock lose rhythmicity in food intake and fermentation product concentration in the cecum, indicating the intricate association between the host circadian rhythms and the microbiota (*65–67*). *Cry1* and *cry2* were upregulated in the ileum for several hours after lactulose treatment, and are known to decrease insulin resistance through the reduction of hepatic gluconeogenesis (*68–71*). Insulin is a hormone that is also associated with satiety, and decreased insulin resistance may have increased satiety in lactulose treated mice (*72*). However, there was no significant change in *cry1/2* in the 3MM liver. It is therefore possible that the changes in expression in the ileum resulted in a signaling mechanism to the liver which resulted in acute changes in glucose metabolism and insulin production.

In conclusion, our work suggests that gut microbiota affects host food intake through the modulation of the circadian clock. We propose a mechanism in which the microbiota breaks down dietary fibers ingested by the host and the metabolic products of microbial metabolism regulate host circadian clock gene expression, which in turn regulates eating behavior. While our results demonstrated an acute effect of diet on the host due to the rapid decoupling of diurnal rhythm and diet, we speculate that the chronic effect of the microbiota on the host through the usual diet of the mouse extends to factors associated with host circadian rhythms such as endocrine signaling, and that the microbiota plays a significant role in maintaining regular satiety-associated behavior through its metabolic byproducts.

## Methods

### Mice

All strains and resources are listed in the key resources table.

Wild type C57BL/6 mice were maintained on a standard diet at the ETH Phenomics center. Mice were re-derived Germ-free or bred with the 3MM microbiota and kept under strict hygienic conditions in breeding isolators. The 3MM microbiota was derived using strains *Bacteroides thetaiotaomicron* (*73*)*, Eubacterium rectale* ATCC, and *Escherichia coli* HS (*74*). Specific pathogen free (SPF) mice were bred and housed in individually ventilated cages. All experiments began with 9-15 weeks of age with mixed-sex groups. Mice were fed autoclaved mouse chow (Kliba Nafag 3807) *ad libitum* for the duration of all experiments and were kept with a dark and light period of 12 hours each. All animal experiments were approved by the Swiss Kantonal authorities (License ZH120/19 and ZH016/21) according to the legal and ethical requirements.

### Metabolic cages

At the start of the experiment, mice were transferred from the breeding isolators to the isolator-housed TSE PhenoMaster (TSE Systems) system and were single housed for the duration of the experiment. Mice were acclimated for 4 days after transfer to the metabolic cages. The TSE PhenoMaster system allows measurements of oxygen, carbon dioxide, and food and water consumption for 160 seconds in 24-minute intervals. Eight metabolic cages were housed in two sterile isolators, and each cage was connected to the system through a HEPA filter. Hydrogen in each cage was measured through a hydrogen sensor (SGP30, Sensirion) which was attached to the TSE Phenomaster system. Air flow in each cage was set to 0.4 L/min. A two-point calibration of all gases using reference gases was performed within 24 hours of each experiment. The TSE Phenomaster data and hydrogen data were combined in the data analysis post-experiment.

### Mouse experiments

On the fifth day after mouse transfer to the TSE Phenomaster, mice were treated by oral gavage with either 132mg Lactulose in 100µL PBS, or 100µL PBS. In cases where 2 lactulose doses were given, the second dose of 132mg was administered 1h after the first. The oral gavage was administered between Zeitgeber 3-4, except when otherwise noted. To measure circadian rhythm changes in the host and changes in the gut, mice were euthanized 5 hours after the gavage. To measure the changes in clock gene expression over 24 hours, mice were euthanized 8 hours, 12 hours, 18 hours, and 24 hours after the gavage. To measure the effect of lactulose on host food intake, mice were euthanized the day following the treatment or two days following treatment. To measure the effect of fermentation product on host food intake, mice were treated between Zeitgeber 3-4 with 66mg of fermentation product in 100µL of PBS, with a 4:3:3 ratio of sodium succinate, sodium acetate, and sodium butyrate, respectively. Mice which did not eat anything in the 24 hours following oral gavage were excluded from the study.

### Bacterial quantification

Mice were euthanized when a peak in hydrogen production was observed following gavage. Wet feces or cecum content were collected from mice and weighed for bacterial quantification. Bacterial DNA was extracted using the DNeasy PowerSoil Pro Kit (Qiagen). qPCR was performed using the FastStart Universal SYBR Green Master Mix (Roche). Primers for each tested strain were diluted to a final concentration of 1µM. DNA was amplified using a QuantStudio 7 Flex instrument (Applied Biosystems) or a StepOnePlus Real-Time PCR System instrument (Applied Biosystems), with an initial denaturation step of 95°C for 10 minutes, followed by 40 cycles of 95°C for 15 seconds and 60°C for 60 seconds. Bacteria were quantified in each run by comparing the obtained Ct values to a standard curve of Ct values for bacterial DNA from a known bacterial cell count in the respective strain, which was extracted using the same protocol. Germ-free cecum content and water samples were used to generate a detection limit. For each experiment, we included a quality control for gnotobiosis, using *Blautia pseudococcoides* as an indicator strain for contaminations. This strain is present in our mouse colony, forms spores, and has been the most likely contaminant in our gnotobiotic sytem. Consequently, probes for *B. pseudococcoides* were included in the qPCR assays, and mice with a C_t_ lower than 26 in these runs were excluded from the study.

### Cecal pH measurement

Cecum content was collected in a 2mL Eppendorf tube (Sarstedt) and snap frozen in liquid nitrogen 5 hours after treatment. At the time of measurement, cecum content was thawed and spun down at 1000rpm for 30 seconds to accumulate all content at the bottom of the tube. A pH sensor (Sevencompact S220, Mettler Toledo) was completely inserted into the cecum content and pH was measured.

### Short chain fatty acid quantification in cecum content

Cecum content was collected in a 2mL Eppendorf tube 5 hours after treatment, snap frozen in liquid nitrogen, and stored at -80°C until further analysis. For UPLC/MS, cecum content was thawed on ice, then homogenized in 70% isopropanol and centrifuged. The supernatant was used for fermentation product quantification using a protocol previously described (*75*). A 7-point calibration curve was generated with fermentation product in known concentrations. Calibrators and samples were spiked with a mixture of isotope-labeled internal standards, derivatized to 3-nitrophenylhydrazones, and then quenched with 0.1% formic acid.

An ACQUITY UPLC system (I-Class, Waters, MA, USA) coupled with an Orbitrap Q-Exactive Plus mass spectrometer (Thermo Scientific) were used for UPLC/MS analysis. To quantify fermentation products, a Kinetex 2.6 µm XB-C18, 50 × 2.1 mm column (Phenomenex) and a flow rate of 250 μL/min was used with a binary mixture of solvent A (water with 0.1% formic acid) and solvent B (acetonitrile with 0.1% formic acid). The gradient starts from 90% of A, then gradually decreases to 75% of A within 6 min and then to 0% of A within 1 min. Then a 0% of A is kept for 1 min and a 90% of A is kept for 1 min to restore the initial solvent ratio. The column was kept at 40°C and the autosampler at 5°C. Acquired raw data were imported to Skyline (V21.1) and MS2 peaks were integrated. A calibration curve for each compound was built based on the ratio between peak area of unlabeled and labeled form using custom R code. Limit of blank was calculated as mean of the concentration detected in technical blank samples ±1.645 standard deviation.

### Short chain fatty acid quantification in medium

All EAM strains were grown to stationary phase in BHI medium supplemented with 1g/L L-cysteine and 2.5g/L hemin. After growth, strains were subcultured into 10mL of Epsilon medium (50mM NaCl, 0.02M NH_4_Cl, 0.028M K_2_HPO_4_, 0.072M KH_2_PO_4_, 50µM MnCl_2_x4H_2_O, 50µM CoCl_2_, 0.4mM MgCl_2_x6H_2_O, 0.5mM CaCl_2_x2H_2_O, 4µM FeSO_4_x7H_2_O, 20mM NaHCO_3_, 5mM L-cysteine, 1.2mg/l hemin, 1mg/l menadione, 2mg/l folinic acid, 2mg/l vitamin B12) with 20mM lactulose as a carbon source and grown overnight. Medium for growing *B. theta* and *E. rectale* was supplemented with 1% tryptone, and medium for growing *E. coli* was not supplemented with tryptone. 1ml of culture was extracted and filtered in a 0.22µm filter, and fermentation product quantification was performed as previously described (*76*). In short, isocratic HPLC was performed in on a Thermo Scientific Ultimate 3000 (Thermo) using 2.5mM H_2_SO_4_ as mobile phase at a 0.4ml/min flow rate. 20µl of sample was injected, separated over a Phenomenex Rezex RoA organic acid H+ (8%) column kept at 40°C, and compounds were detected using a refractive index detector. Data was recorded for 40min after injection for each sample. Data was analyzed using Python and the package hplc-py (*77*).

### Determining water content of feces and cecum content and total fecal output

Fresh fecal pellets and cecum content were collected 5 hours after treatment and the wet weight was measured. Samples were then lyophilized (Alpha 2-4 LD plus, Martin Christ GmbH) for 48 hours and dry weight was measured. The ratio of dry weight to wet weight was calculated to estimate the water content in the feces and cecum content after the gavage.

For measuring total fecal output after gavage, the bedding in each cage was replaced at the time of the oral gavage. At the end of the experiment, all fecal pellets were collected from the bedding in each cage, and lyophilized to determine the total fecal dry weight that was produced since lactulose gavage. This dry weight of the feces was measured and was then normalized by the water content in fresh feces collected from each mouse to estimate total fecal wet weight produced per animal in the respective time window. For this measurement, we disregard coprophagia, which might lead to a slight underestimation of total fecal output.

### RNA extraction

Mouse ileum, cecum, and colon tissues were collected in RNAlater (Thermo Scientific) and snap frozen in liquid nitrogen. Samples were stored at -80°C until further analysis. For RNA extractions, samples were thawed on ice and weighed. RNA was extracted using a modified protocol of the Direct-zol RNA kit (Zymo). Samples were mixed with Trizol (Thermo Scientific) and incubated for 5min at room temperature. Samples were then homogenized twice with 0.1mm beads (BioSpec Products) and 1.0mm silicon carbide beads (BioSpec Products), for 90s at 30Hz. Further Trizol was mixed with the samples, and centrifuged. Supernatant was mixed with chloroform (Sigma Aldrich) and centrifuged. Supernatant was then washed in 100% ethanol and centrifuged in collection tubes. RNA was washed in Pre-wash buffer and incubated for 15min at room temperature with 60 units of DNAse. Finally, RNA was washed in Wash buffer and eluted in Nuclease-free water (Invitrogen).

RNA concentration and quality were measured using a nanodrop machine (ND-1000 Spectrophotometer, Thermo Scientific) and RNA was diluted to a final concentration of 20ng/µL in nuclease-free water. Reverse transcription was done using the High-Capacity cDNA Reverse Transcription Kit (Applied Biosystems). cDNA was diluted 1:25 in nuclease-free water for qPCR.

### Circadian clock gene expression

Host circadian clock gene primers were used to estimate expression using RT-PCR. A housekeeping gene (rpl19) was used for normalization. The RT-PCR was carried out as previously described CITE, and sample extracted RNA was included in a reaction with the housekeeping gene to control for DNA contamination.

### GLP-1 and PYY measurements

Mice were anaesthetized with isofluorane before exsanguination and tissue sampling. Portal vein blood was collected. Plasma was isolated and immediately transferred to tubes with 20µL of DPP IV inhibitor (Sigma Aldrich) and immediately flash frozen. GLP-1 and PYY measurements were done using an ELISA kit (Crystal Chem), according to the manufacturer’s instructions.

### Leptin and Ghrelin measurements

Mice were euthanized using CO_2_. Blood was collected by cardiac puncture, and serum was collected and flash frozen. Leptin and total Ghrelin measurements were done using an ELISA kit (Sigma Aldrich), according to the manufacturer’s instructions

## Data analysis

### Phenomaster quality control

For all measurements, a variable of zeitgeber time (h) was created where 0-12 represents the light phase and 12-24 represents the dark phase animals experience in the animal facility. The day number of the experiment was calculated based on the zeitgeber time of the first measurement. Each new day began at zeitgeber time 0, at the start of the light cycle. To account for faulty measurements due to disruptive events and measurement noise, some datapoints were excluded from the raw dataset: food intake above 1g and water intake above 1ml in one 24-minute interval for one mouse was considered to have been caused by human disruption and was excluded, food intake of 0.01g as well as negative food and drink values were considered measurement noise and were excluded; for each mouse, the interquartile range of food and water intake measurements was calculated, and values greater than the 75^th^ percentile + 1.5 times the interquartile range were considered outliers and excluded from the dataset. Potential sources of outlier measurements include food and water loss during handling of the experimental setup, as well as leaky water bottles. We analyzed whether the probabilities for excluding values was the same between the experiments show in in Figures 4A/S5A, 4B/S5B, and S5F, and found that they were significantly different (Chi-square test, p=3.9 x 10^-18^), which could be due to differences between the experiments in the number of times cages were handled. Exclusion probabilities between treatment and control were significantly different in Figure 4A/S5A (Fisher’s exact test, p= 1.7 x 10^-6^), but not significantly different in the experiments shown in Figures 4B/S5E and S5F (Fisher’s exact test, p=0.71 and p=0.10, respectively). While we cannot strictly rule out any influence of our data exclusion strategy, these results speak against a systematic effect of treatment on behavior that would lead to changes the amount of excluded data points. After data cleanup, mice which had a total daily food intake difference of more than 1g in the two days prior to treatment were considered not acclimated to the metabolic cages and were excluded from further analysis.

### Hydrogen sensor data

The hydrogen sensors attached to the TSE Phenomaster system (Sensirion SGP30) measured hydrogen every second, and hydrogen measurements were recorded in a separate file. The median and maximum hydrogen values for each experimental box and reference box measurement were calculated. Median hydrogen values were used for further analysis. If the reference box measurements remained visually constant for the duration of the experiment, the hydrogen data was accepted and merged with the Phenomaster data for further analysis.

### Metabolic cage statistical analysis

For estimating daily mouse food intake and hydrogen production, data was extracted from the raw dataset in the two days prior to mouse treatment. For each mouse, the average hydrogen value and food intake measurement per timepoint was calculated. A general additive model was used to fit a curve (using R function mgcv::gam) to these points using the formula y∼s(x; bs=”cs).

For estimating food intake per day, data was extracted from the beginning of the day prior to mouse treatment to the end of the first dark cycle following treatment, resulting in data from two dark cycles. Cumulative food intake and total food intake were calculated per mouse for each dark cycle, and food intake rate was calculated as the slope of the linear regression of cumulative food intake. A paired T-test was used to compare food intake rate and total food intake for each mouse in the dark cycle prior to treatment and in the dark cycle following treatment.

### Energy harvesting estimations

For estimating the indirect caloric intake of the EAM mouse in the lactulose treatment, we made a deliberate overestimation, assuming that the entire energy contained in the administered lactulose would reach the mice. We estimated the combustion enthalpy of lactulose to be the same as that of lactose (1346 kcal/mol), and the combustion enthalpy of 132mg (0.39mmol) of lactulose is therefore 0.52kcal (*78*). For estimating the caloric intake of the EAM mouse in the fermentation product treatment, it was assumed that 100% of fermentation product given to the host were used for energy harvesting. The combustion enthalpies and masses provided to the host of acetate (0.21kcal/mmol, 19.8mg), butyrate (0.52kcal/mmol, 19.8mg), and succinate (0.36kcal/mmol, 26.4mg) were used to calculate the maximum possible calories provided to the host (0.19 kcal).

Energy density of mouse chow was previously measured to be 3.94kcal/g [35]. We calculated the total caloric intake per mouse for the dark cycle preceding and following lactulose treatment based on the energy from food intake and indirect feeding from the microbiota and fermentation product treatment. A paired t-test was used to compare caloric intake for each mouse in the dark cycle prior to treatment and in the dark cycle following treatment.

### Gene expression

A fluorescence threshold of 0.3 was set for all primers. For data analysis, the ΔΔC_t_ method was used to estimate relative gene expression for each experiment (*79*). In all samples, the C_t_ of the gene of interest was first normalized by that for the housekeeping gene rpl19. For samples collected over 24 hours (data shown in Figure 3C), gene expression values were then normalized to those of untreated mice euthanized at 9AM. For all other samples, gene expression was normalized to the PBS-treated control mice.

For estimating clock gene expression differences in gastrointestinal tissue samples, and for calculating fermentation product concentration in cecum content, unpaired t-tests were used to compare both groups. Benjamini Hochberg multiple testing correction was used on both these datasets. For all further comparisons between treatment groups, unpaired t-tests were used. All data analysis was carried out using R V4.2.0.

## Acknowledgements

We would like to thank Daniel Hoces for providing the base R script which was adapted for analyzing the Phenomaster data. We would further like to thank Sven Nowok and Dominik Bacovcin and the animal caretakers at the EPIC facility at ETHZ for maintaining the mouse lines. We would like to thank Sandra Kaiser, Bidong Nguyen, Sanne Kroon, and Wolf Dietrich-Hardt for providing the Lactulose dosage for this study.

## Author contributions

GG, ES, and MA designed the research. GG, MA, CM, SO, AWO, ECB, JL, and DK performed experiments. GG and MA evaluated and analyzed the data and wrote the manuscript in cooperation with all authors.

## Lead contact

Further information and requests for resources and reagents should be directed to and will be fulfilled by Markus Arnoldini or Giorgia Greter.

## Materials availability

All materials are available upon request to the corresponding author.

## Data and code availability

All data relevant for generating figures is available in a permanent data repository (doi: 10.5281/zenodo.20624460).

## Supporting information

**Supplementary Figure 1.**
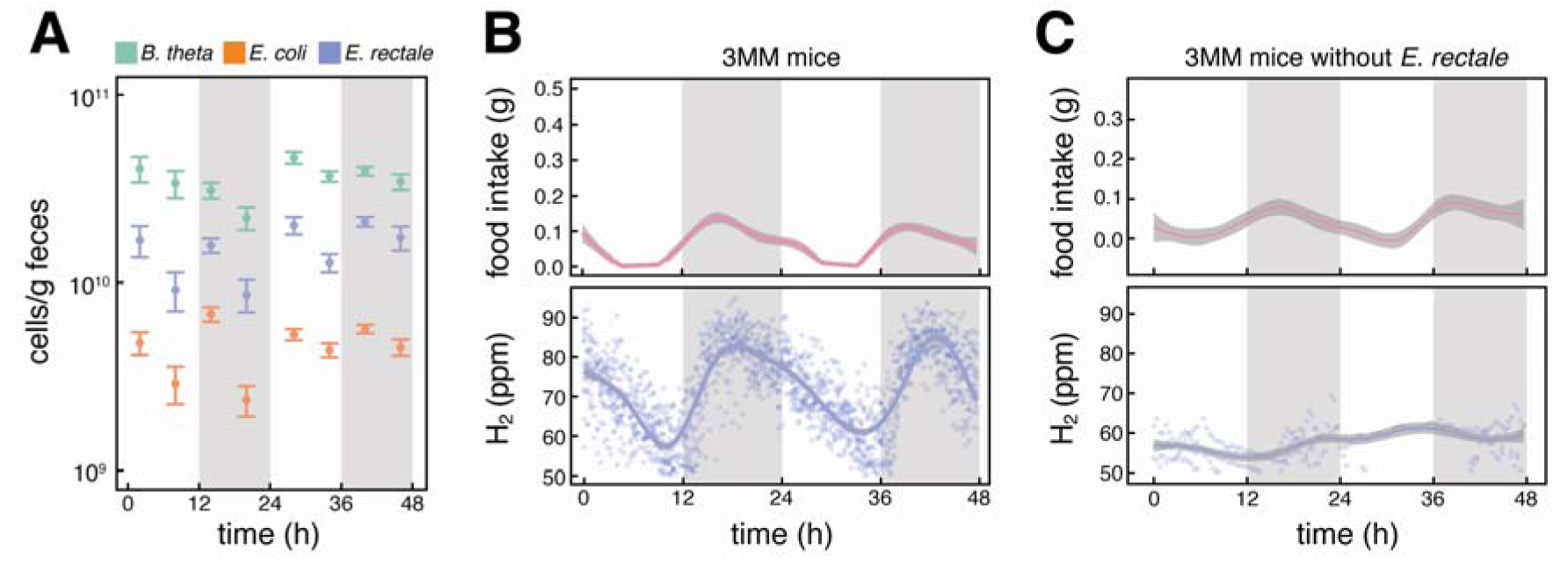
*E. rectale* is the principal hydrogen producer in the 3MM mirobiota. **(A)** Microbial cell counts based on qPCR in 3MM mouse feces. Fecal pellets were sampled over 2 days in 8 3MM mice. **(B)** Food intake and hydrogen production in 3MM mice (n=16) over two days, with measurements taken every 24min. Trend lines were plotted using the geom_smooth function in the ggplot2 package in R with standard settings, shaded areas around the trend lines represent standard error of the mean. **(C)** Food intake and hydrogen production of gnotobiotc mice colonized with *B. theta* and *E. coli* only (n=4). In all panels, shaded background indicates dark periods in the animal facility, from 6pm (zt=12) to 6am (zt=24).

**Supplementary Figure 2.**
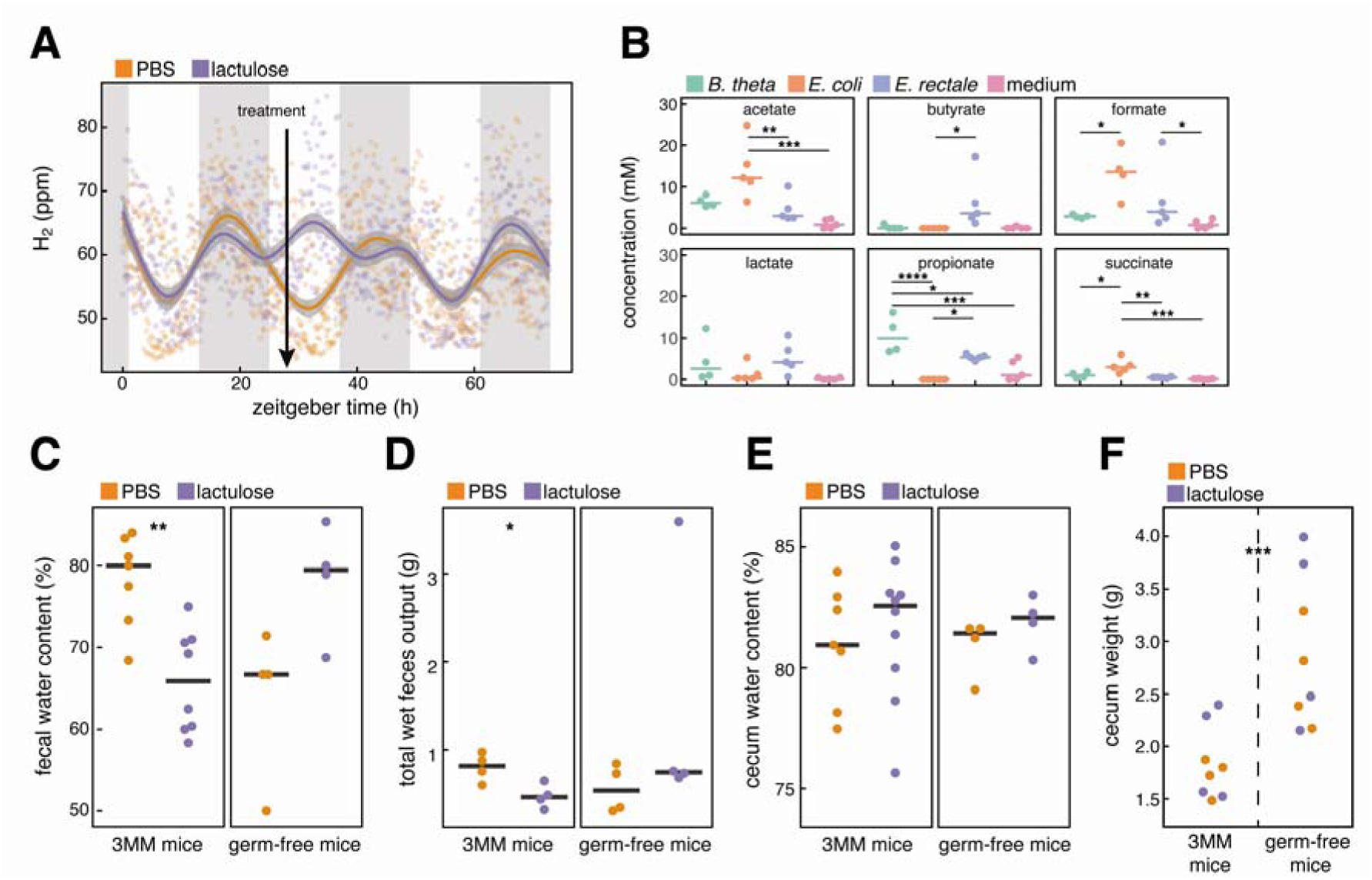
Lactulose changes the diurnal pattern of microbial metabolism. **(A)** Hydrogen production temporarily increases after treatment with lactulose (n=5), but not after treatment with PBS (n=6), in SPF mice. Treatment is indicated by a black arrow. **(B)** Production of fermentation products by the bacterial strains constituting the 3MM microbiota grown in axenic culture in media with lactulose as carbohydrate source. Measurements were taken in stationary phase (*, **, ***, ****, indicate p<0.05, 0.01, 0.001, and 0.0001, respectively; ANOVA with Tukey’s post-hoc test). **(C)** Fecal water content in 3MM mice treated with lactulose (n=8) and PBS (n=7), and germ-free mice treated with lactulose (n=4) and PBS (n=4). The water content decreases significantly upon lactulose treatment in 3MM mice (p<0.01, unpaired t-test). **(D)** Estimated production of total fecal wet weight during 5h in 3MM mice after treatment with lactulose (n=4) and PBS (n=4), and in germ-free mice treated with lactulose (n=4) and PBS (n=4). This data was calculated by experimentally measuring total fecal dry weight and correcting it with the fecal water content determined for the respective microbiota and treatment in **(C)**. Output of fecal wet weight increases significantly in 3MM mice after lactulose treatment (unpaired t-test, p<0.05). **(E)** Water content of cecum content in 3MM mice treated with lactulose (n=10) and PBS (n=7). **(F)** Cecum weight of 3MM and germ-free mice treated with PBS or lactulose. Germ-free mice have significantly higher cecum weights than 3MM mice (p<0.01, unpaired t-test), which already have higher cecum weights than conventionally colonized mice (around 0.5g)(*35*).

**Supplementary Figure 3.**
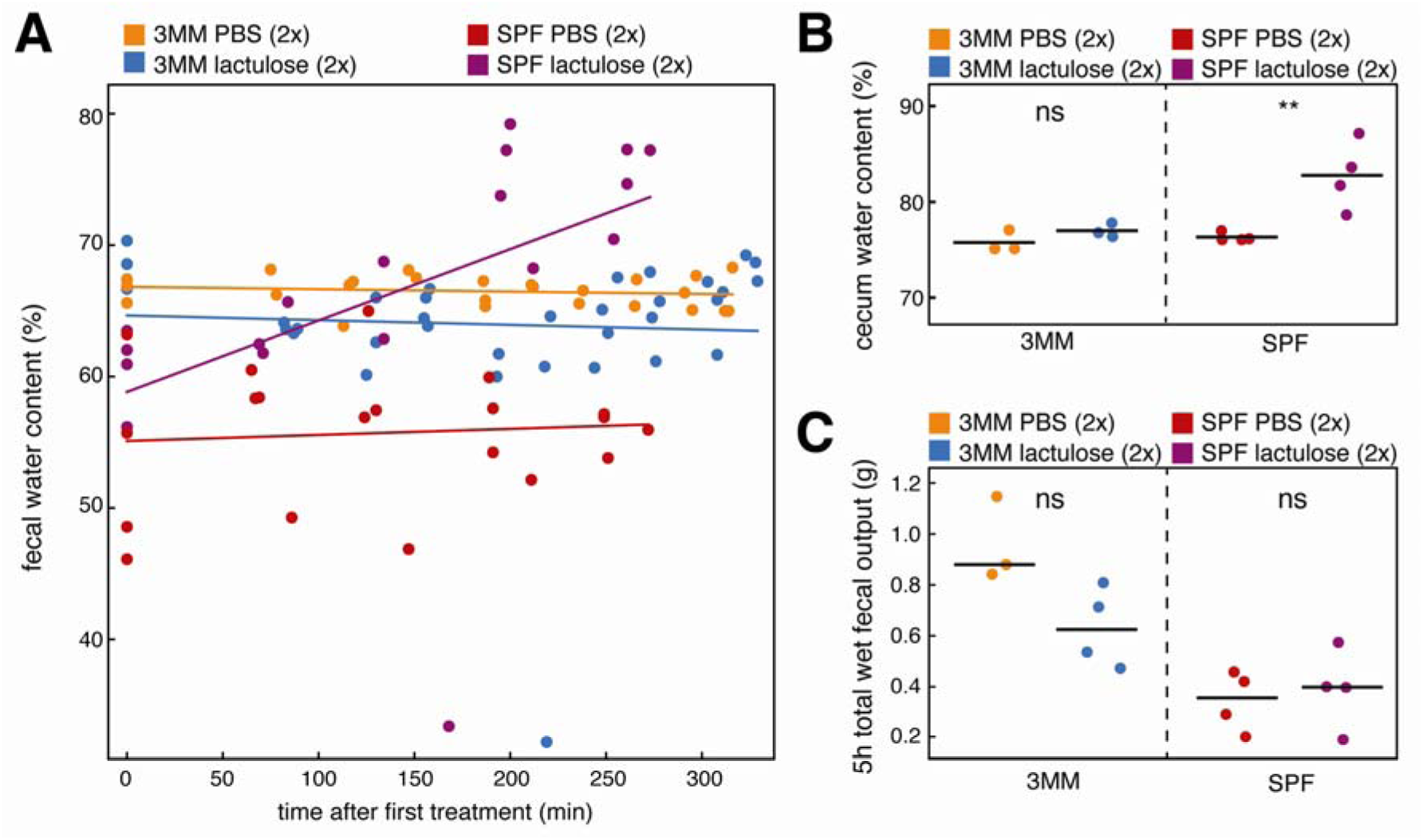
Even higher dose lactulose treatment does not lead to an osmotic effect in 3MM mice, but does so in SPF mice. **(A)** Water content in fresh fecal pellets collected in regular intervals after PBS or lactulose treatment, in 3MM and SPF mice. Lines indicate linear regression on the data. Fecal water content does not increase after PBS treatment in 3MM or SPF mice. Lactulose treatment does also not lead to an increase in water content in 3MM mice, but did lead to an increase in SPF mice, indicating that the osmotic effect of lactulose depends on the colonization state of the animals. **(B)** Cecum water content 5h after treatment. Only lactulose treated SPF mice showed a significant increase in water content in cecum content 5h after treatment (p<0.01, ANOVA with Tukey’s post-hoc test). **(C)** Total fecal output during 5h after treatment was not significantly different between groups (ANOVA). This data was collected the same way as in Figure S2D, by collecting all fecal pellets from cage bedding, measuring their dry weight, and correcting it by the mean water content of the fresh feces in the respective group.

**Supplementary Figure 4.**
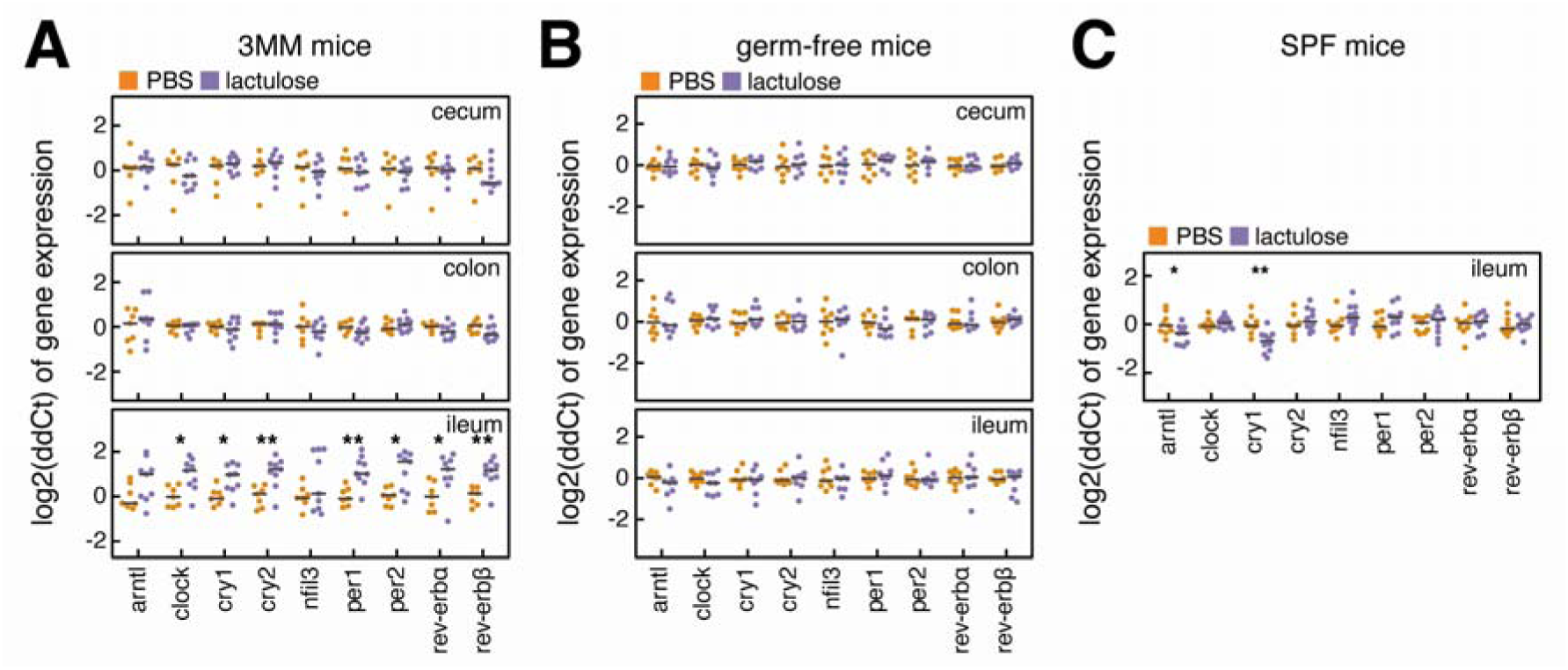
Targeted induction of microbial metabolism by lactulose treatment acutely alters clock gene expression in small intestinal tissue. **(A)** Clock gene expression measured by qPCR in tissue samples of ileum, cecum, and colon of 3MM mice after treatment with lactulose (n=10) and PBS (n=7). Significant differences in gene expression between groups were detected in samples from the ileum (unpaired t-test, multiple testing correction). **(B)** Clock gene expression in tissue samples of ileum, cecum, and colon of germ-free mice after lactulose (n=8) and PBS (n=8) treatment. No significant differences were found in gene expression levels between treatment groups (unpaired t-test, multiple testing correction). **(C)** Clock gene expression in ileum tissue of SPF mice after treatment with lactulose (n=5) or PBS (n=5). Cry1 expression was significantly different between treatment groups (unpaired t-test, Benjamini-Hochberg correction). In all panels, * and ** indicate p<0.05 and p<0.01, respectively.

**Supplementary Figure 5.**
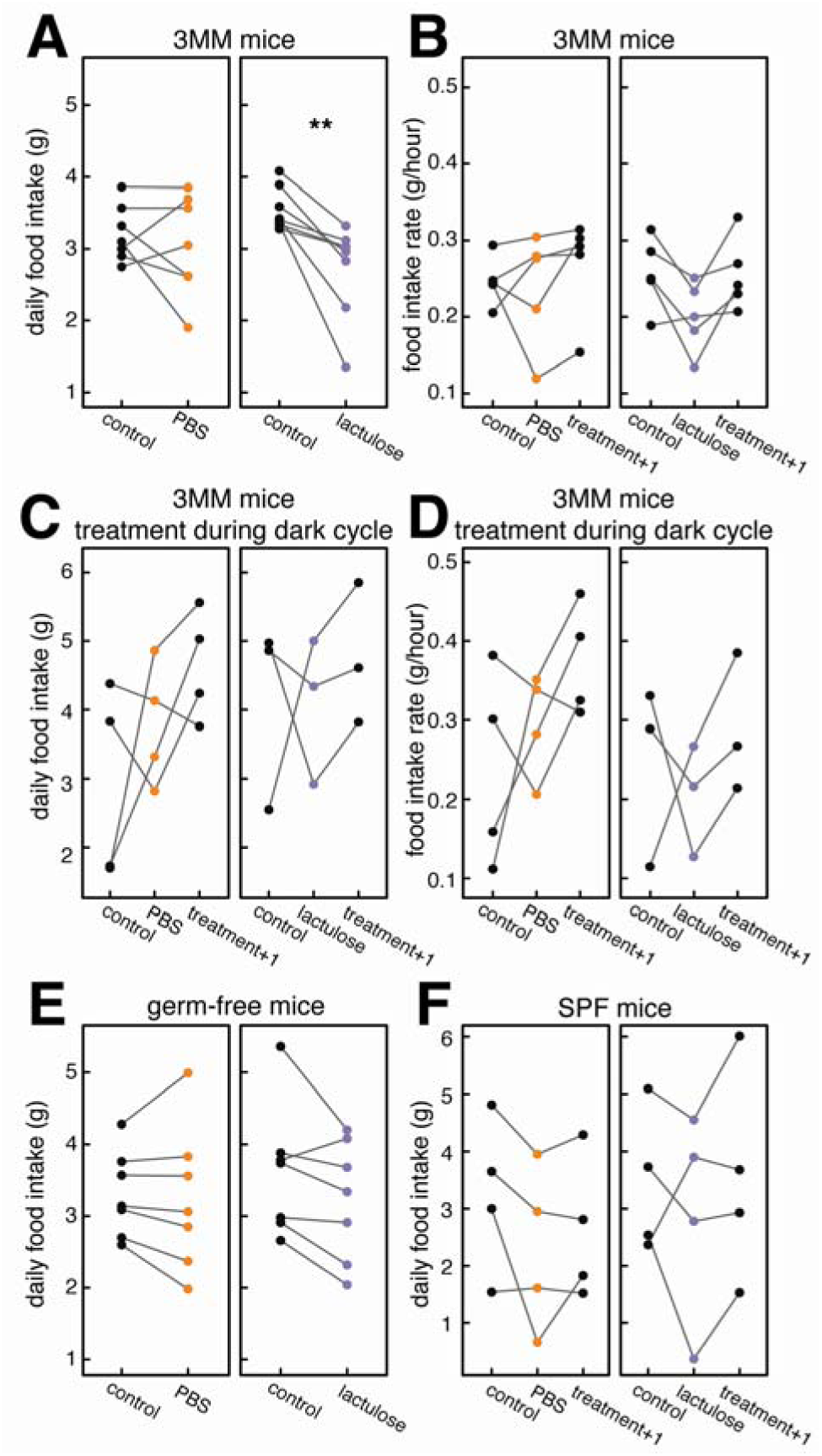
Induction of microbial metabolism transiently decreases host food intake in the following active cycle. **(A)** Total food intake in 3MM mice before treatment (control) and after lactulose (n=8) and PBS (n=7) treatments. Food intake was significantly lower after lactulose treatment (paired t-test, p<0.01). **(B)** Food intake rate in 3MM mice in the dark phase before treatment (control) and the two dark phases following treatment (treatment, treatment+1). The food intake rate is not significantly different between the days (ANOVA). **(C)** Total food intake in 3MM mice before treatment (control) and in the two dark cycles after treatment with lactulose (n=3) and PBS (n=4), when treated at 8:30pm (zt=14.5, 2.5h into the dark cycle). No significant differences were found between groups (ANOVA). **(D)** Food intake rate for the experiment shown in in **(C)**. **(E)** Total food intake in germ-free mice in the dark cycle before treatment (control) and the dark cycle after treatment with lactulose (n=7) and PBS (n=7). No significant differences were found between the days (paired t-test). **(F)** Total food intake of SPF mice before treatment (control) and in the two dark cycles after treatment with lactulose (n=4) and PBS (n=4). No significant differences were found between the days (ANOVA).

**Supplementary Figure 6.**
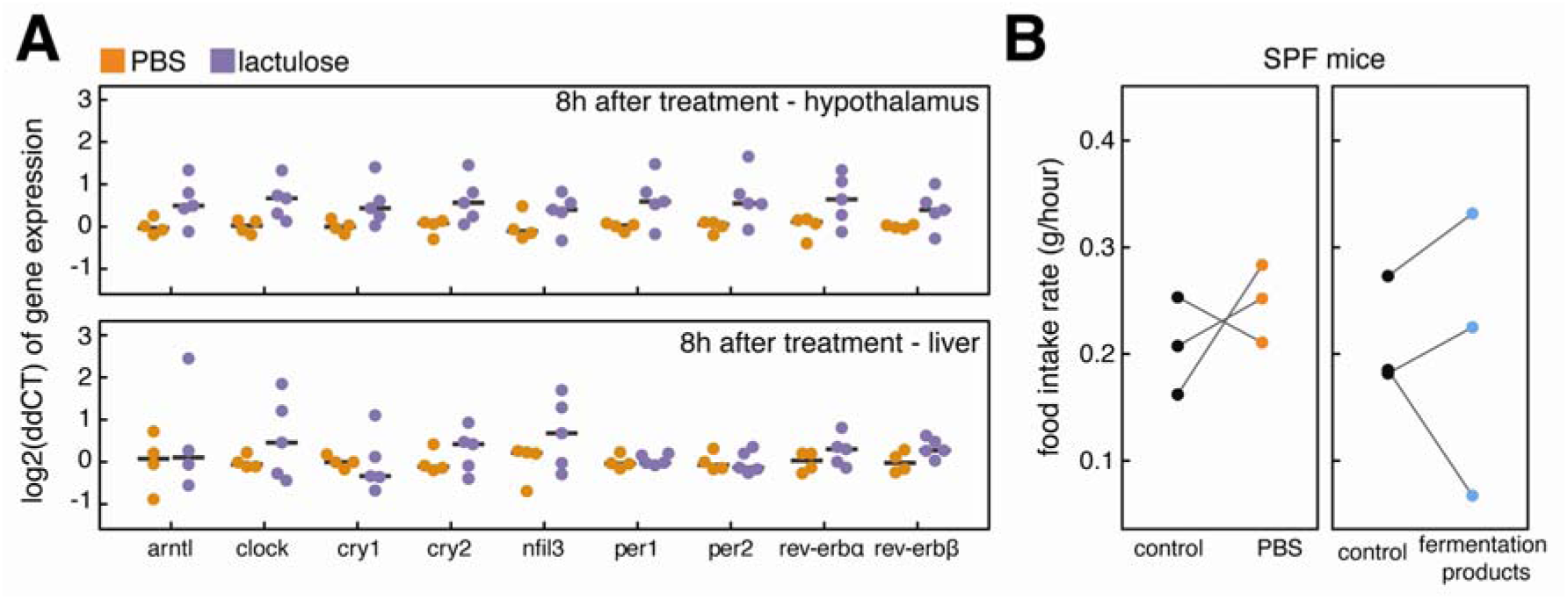
Clock gene expressing in perpheral tissues and food intake rates in SPF mice after PBS or lactulose treatment. **(A)** Clock gene expression in hypothalamus and liver of EAM mice after lactulose (n=5) and PBS (n=4) treatment, measured via qPCR. Gene expression levels are not significantly different between groups (unpaired t-test). **(B)** Food intake rates of SPF mice in the dark phase before treatment (control) and the dark phase following treatment with fermentation products. Food intake rates did not differ significantly between the days (paired t-test).

**Supplementary Figure 7.**
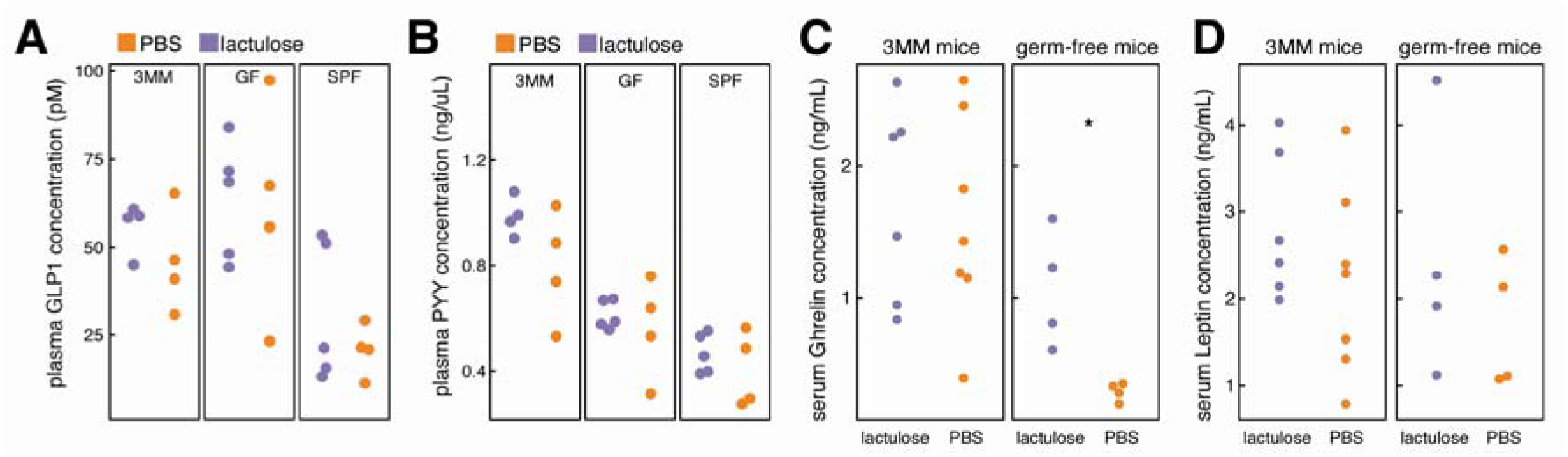
Concentration measurements of metabolic hormones after treatment with PBS or lactulose. **(A)** Total GLP1 concentration in portal vein plasma 5h after lactulose or PBS treatment in 3MM, germ-free, and SPF mice. We found no significant differences between groups (unpaired t-test). **(B)** PYY concentration in portal vein plasma after lactulose or PBS treatment in 3MM, germ-free, and SPF mice. Groups were not significantly different (unpaired t-test). **(C)** Total Ghrelin concentration in serum from cardiac puncture after lactulose or PBS treatment in 3MM and germ-free mice. No significant differences were found between groups in 3MM mice. Ghrelin was significantly increased (p<0.05) in germ-free mice after lactulose treatment (unpaired t-test). **(D)** Total Leptin concentration in serum from cardiac puncture after lactulose or PBS treatment in 3MM and germ-free mice. Concentrations were not significantly different between groups (unpaired t-test).

**Table S1.**
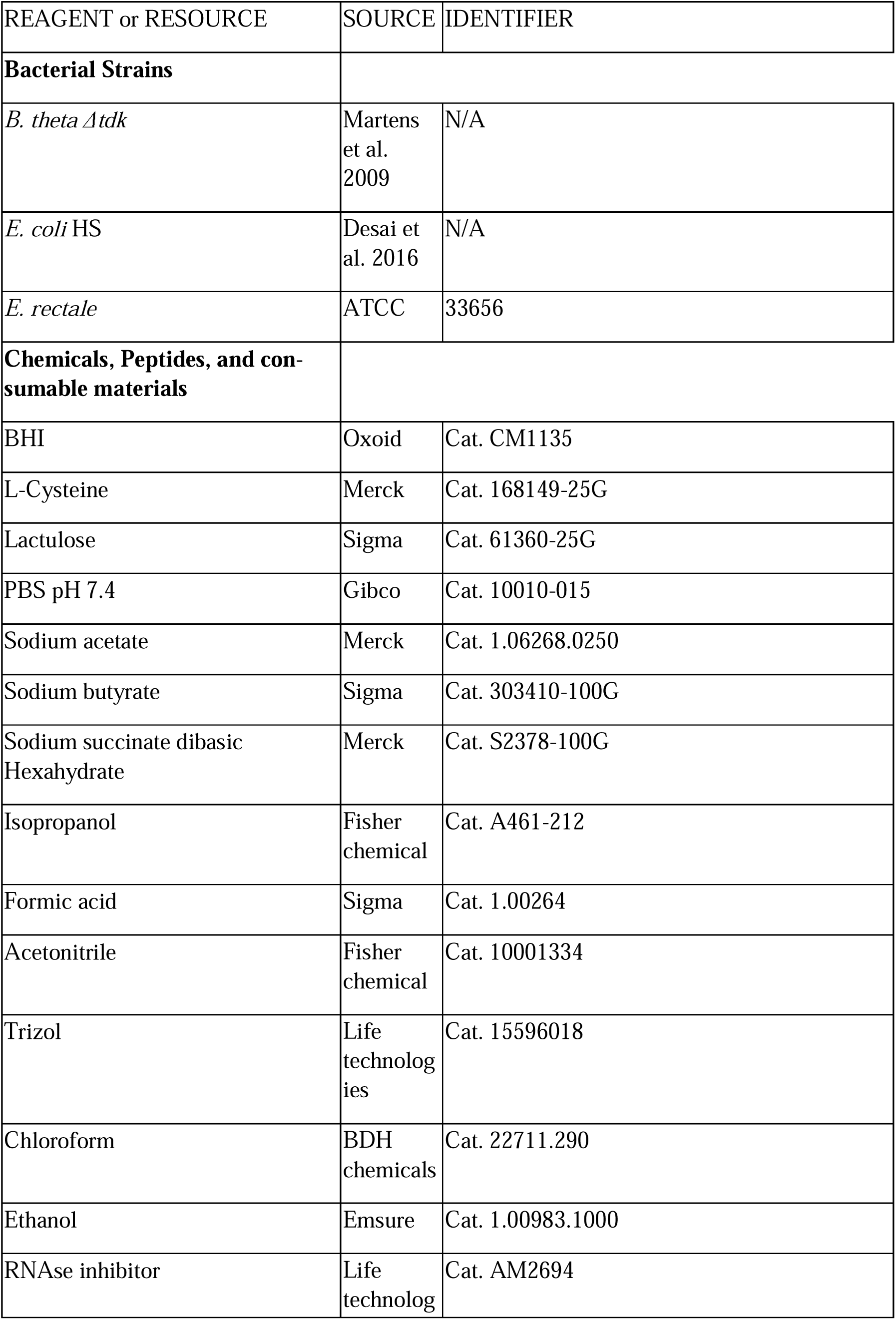
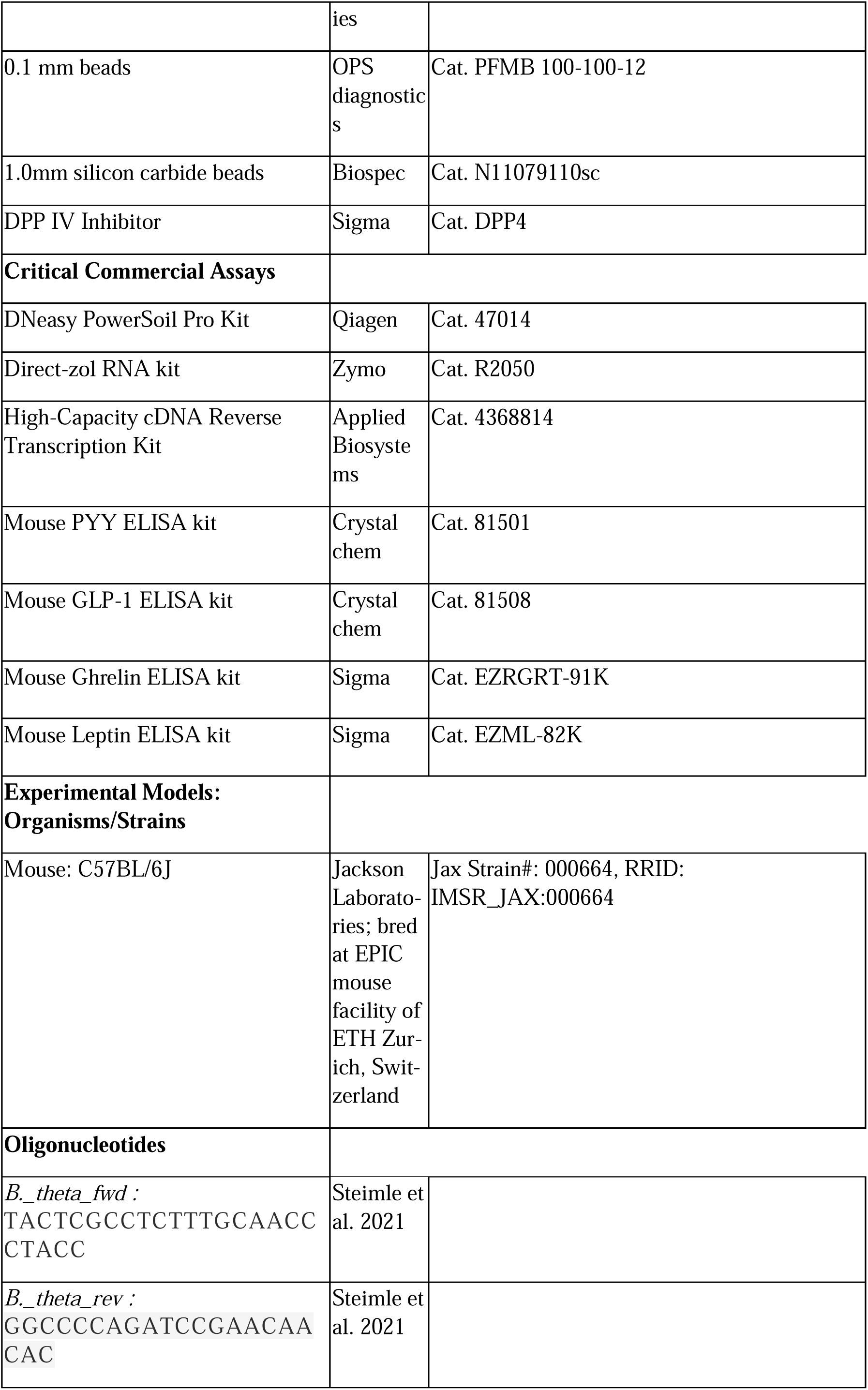
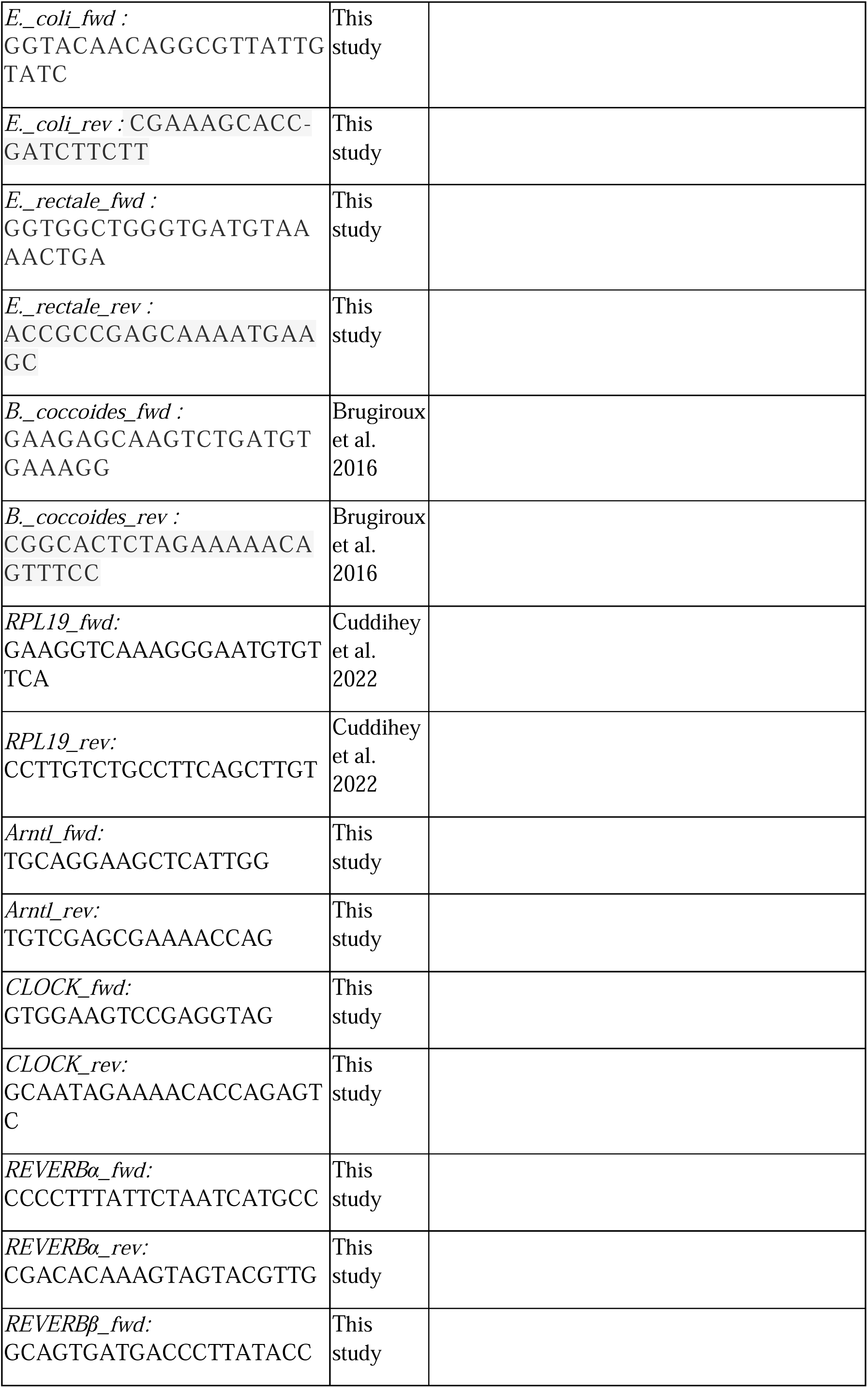
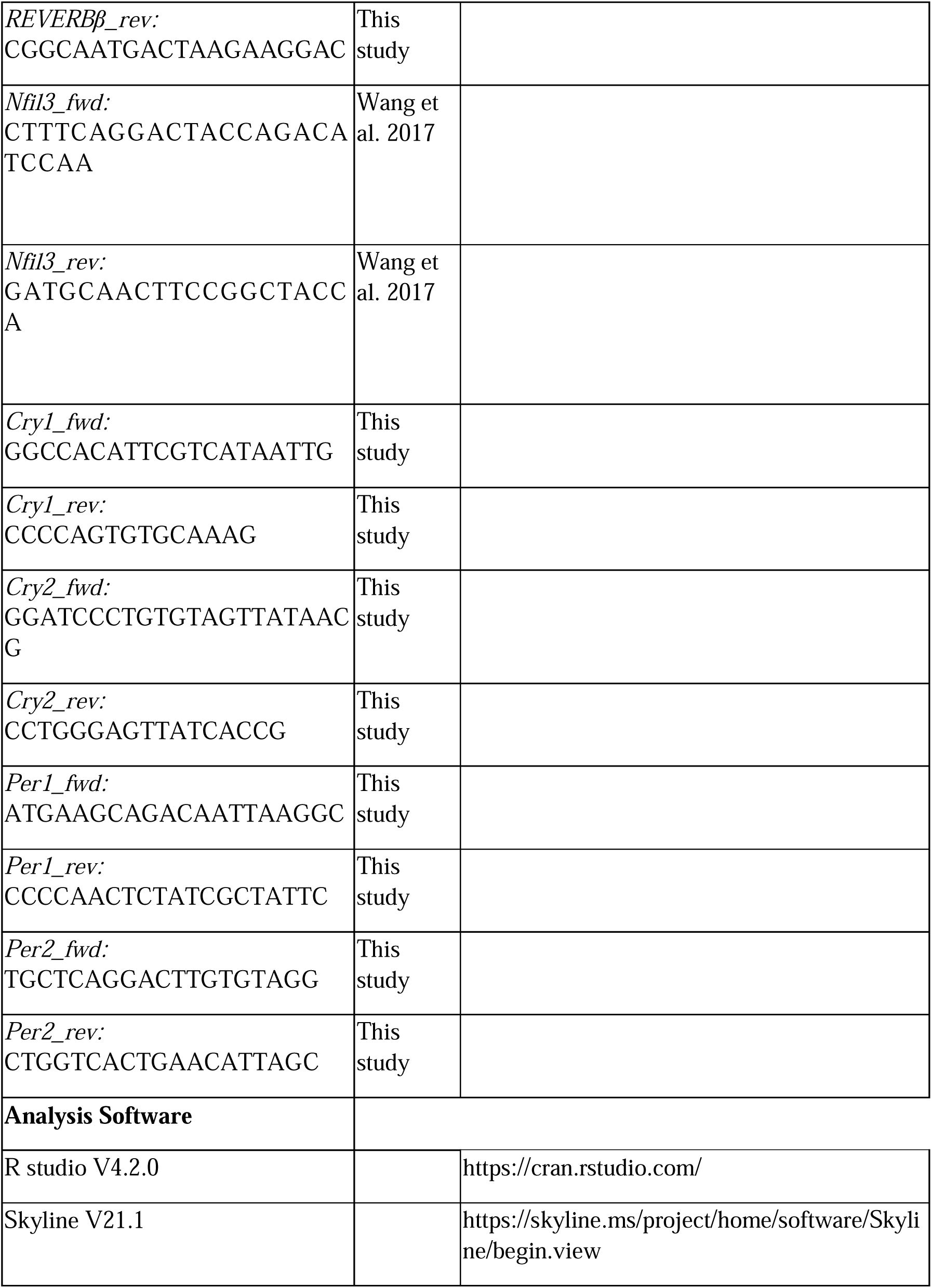
Key resource table.

